# Membrane Diffusion Occurs by a Continuous-Time Random Walk Sustained by Vesicular Trafficking

**DOI:** 10.1101/208967

**Authors:** Maria Goiko, John R. de Bruyn, Bryan Heit

## Abstract

Diffusion in cellular membranes is regulated by processes which occur over a range of spatial and temporal scales. These processes include membrane fluidity, inter-protein and inter-lipid interactions, interactions with membrane microdomains, interactions with the underlying cytoskeleton, and cellular processes which result in net membrane movement. The complex, non-Brownian diffusion that results from these processes has been difficult to characterize, and moreover, the impact of factors such as membrane recycling on membrane diffusion remains largely unexplored. We have used a careful statistical analysis of single-particle tracking data of the single-pass plasma membrane protein CD93 to show that the diffusion of this protein is well-described by a continuous-time random walk in parallel with an aging process mediated by membrane corrals. The overall result is an evolution in the diffusion of CD93: proteins initially diffuse freely on the cell surface, but over time, become increasingly trapped within diffusion-limiting membrane corrals. Stable populations of freely diffusing and corralled CD93 are maintained by an endocytic/exocytic process in which corralled CD93 is selectively endocytosed, while freely diffusing CD93 is replenished by exocytosis of newly synthesized and recycled CD93. This trafficking not only maintained CD93 diffusivity, but also maintained the heterogeneous distribution of CD93 in the plasma membrane. These results provide insight into the nature of the biological and biophysical processes that can lead to significantly non-Brownian diffusion of membrane proteins, and demonstrate that ongoing membrane recycling is critical to maintaining steady-state diffusion and distribution of proteins in the plasma membrane.

## Introduction

The plasma membrane is a complex organelle in which dynamic structural and organizational features enable processes such as receptor-mediated signaling, endocytosis and exocytosis, and intercellular interactions. Many of these processes rely on self-organizing molecular complexes whose assembly, composition, and lifetime are dictated by the diffusional behaviors of their constituent members. Membrane diffusion is a complex biophysical process which differs significantly from free Brownian diffusion (reviewed in 1, 2). As a result, the apparent diffusional behavior of proteins in cellular membranes determined from experimental data depends on factors such as the sampling rate and the duration of the measurement. At high temporal resolution, some membrane proteins have been shown to undergo free Brownian motion over length scales of < 200 nm (3), although this scale is molecule and cell-dependent. Over longer times their motion becomes non-Brownian, and molecules instead undergo hop-diffusion characterized by transient periods of trapping within confinement zones separated by “hops” from one confinement zone to another (3–5). This “wait-and-jump” behavior is reminiscent of a continuous-time random-walk (CTRW), a form of anomalous diffusion characterized by particles which move by a series of “jumps” separated by waiting periods of random duration (6, 7). Further deviations from Brownian diffusion may result from molecular crowding, inter-protein interactions and interactions of diffusing proteins with membrane microdomains (8–10). Despite the complex non-Brownian motion of individual molecules, techniques such as fluorescence recovery after photobleaching, which analyze diffusion over second-to-minute time scales, show Brownian-like diffusion, albeit with effective diffusion coefficients one to two orders of magnitude lower than those observed in model membranes that are free of confinement zones and have a low protein density (9).

This apparent change in diffusional behavior under different observational conditions is thought to be an emergent property reflecting the influence of multiple processes occurring over a range of time and length scales. Brownian diffusion of both proteins and lipids has been reported in simple model membranes (11, 12), and over short periods of time (nano-to-milliseconds) in cellular membranes (3). Molecular modeling of protein rich membranes suggest that Brownian-like diffusion of proteins should give way to anomalous diffusion at time-scales longer than a few microseconds (13), while in cellular membranes the transition from Brownian to anomalous hop-diffusion has been observed at time scales of a few tens of milliseconds and at spatial scales between 20 and 200 nm (14). The disagreement between these model and observational studies may be due cellular factors unaccounted for by current models, or by the current spatial (10-20 nm) and temporal (hundreds of microseconds) limitations of single molecule tracking systems. Hop diffusion results from interactions between diffusing molecules and barriers comprised of membrane-proximal actin filament “fences”, attached to “pickets” of actin-bound transmembrane proteins which have been observed to form a percolation barrier in the membrane (15, 16); similar barriers comprised of the cytoskeletal proteins spectrin and septins have also been reported (17, 18). These picket-fence structures create confinement zones, termed “corrals”, ranging from 40 to ~400 nm in size, which restrict the diffusion of transmembrane proteins, lipids, and lipid-anchored proteins on both the cytosolic and extracellular leaflet of the plasma membrane. Diffusing proteins and lipids are transiently trapped within corrals, then “hop” between corrals either by successfully diffusing through the percolation barrier, or by diffusing through gaps created in the “picket fence” by actin turnover (4, 5). Diffusion of proteins is further modified by interactions with membrane microdomains such as lipid rafts, and with other membrane, cytosolic and extracellular proteins (10, 15, 19–24). These interactions take place on timescales ranging from sub-millisecond (protein-protein and protein-lipid interactions), to tens of milliseconds (rafts), to seconds (actin corrals), to hours or even days (focal contacts). A final layer of complexity is added by the cellular processes of exocytosis and endocytosis, which respectively deliver and remove material from the plasma membrane. These processes also occur over a wide range of time scales, from a few tens of milliseconds during ultra-rapid membrane recycling in neurons, to several minutes for recycling endosome pathways (25, 26).

A consequence of this complex environment is that measurements of diffusional behavior are highly dependent on the temporal resolution and duration of the experiments used to observe diffusion. This has made it difficult to develop a mathematical framework which realistically describes diffusion within the plasma membrane of eukaryotic cells (2, 27). Moreover, diffusion is conventionally quantified using time-averaged mean-squared displacement (TAMSD), which measures changes in molecular position as a function of the lag time between observations, without regard to when in real-time those observations were made. While this approach is statistically robust and sensitive to processes which operate on different time-scales, it obscures processes which evolve over real-time. Of greater concern is that some methods used to quantify diffusion in biological systems assume that cellular diffusion is scale-invariant (i.e., that diffusion measured over one temporal or spatial range will be identical to diffusion measured over different temporal or spatial ranges), ergodic (i.e., that every region of space is visited with equal probability), or occurs in a homogeneous environment – assumptions which recent measurements of cellular diffusion clearly demonstrate are untrue (28–30).

Several models have been developed to describe non-Brownian diffusion. These include fractional Brownian motion (FBM), CTRWs, and motion in a fractal environment, all of which have been applied to describe the anomalous diffusion of proteins in a biological context (2, 29, 31). Brownian motion, FBM and diffusion in fractals are scale invariant and ergodic. In Brownian motion the TAMSD as a function of lag time τ is equivalent to the ensemble-averaged MSD (EA-MSD) as a function of measurement (real) time *t*. For non-Brownian systems, this equivalence may not hold. Different models predict different relationships between these quantities (reviewed in 31). For example, simple Brownian diffusion is accurately modeled by a random walk, in which a particle moves a fixed distance in a random direction at each time step. In a CTRW, in contrast, the particle is ``trapped’’ for a random waiting time before making each jump. The former process is ergodic, while, if the waiting times are power-law distributed, the latter is not. In principal, the two models can be distinguished from each other by the lag-time and measurement-time dependence of the time- and ensemble-averaged MSDs.

In this study, we carry out a careful statistical analysis of long-duration single-particle tracking data to characterize the anomalous diffusion of the type-I transmembrane protein CD93, a group XIV C-type lectin linked to the plasma membrane by a single transmembrane domain and bearing a short intracellular domain. While CD93’s ligands and signaling pathways remain to be elucidated, it has established roles in efferocytosis, angiogenesis, and cell adhesion (33–35). Our analysis demonstrates that the diffusion of CD93 is well-described by a continuous-time random walk model, evolves over real-time, and is sustained by endocytosis and exocytosis.

## Materials and Methods

### Materials

The 12CA5 hybridoma was a gift from Dr. Joe Mymryk (University of Western Ontario). Lympholyte-poly, CHO cells and Cy3-conjugated rabbit-anti-mouse Fab fragments were purchased from Cedarlane Labs (Burlington, Canada). All tissue culture media, Trypsin-EDTA 100X antibiotic-antimycotic solution and fetal bovine serum (FBS) were purchased from Wisent (St. Bruno, Canada). 18mm, #1.5 thickness round coverslips were purchased from Electron Microscopy Sciences (Hatfield, PA). Anti-human CD93 was purchased from eBioscience (Toronto, Canada). Chamlide magnetic Leiden chambers were purchased from Quorum Technologies (Puslinch, Canada). GenJet Plus *In Vitro* DNA Transfection Reagent was purchased from Frogga Bio (Toronto, Canada). Leica DMI6000B microscope, objectives and all accessories were purchased from Leica Microsystems (Concord, Canada). Computer workstation was purchased from Stronghold Services (London, Canada). Matlab equipped with the parallel processing, statistics, image processing and optimization toolboxes was purchased from Mathworks (Natick, MA). Prism statistical software was purchased from Graphpad (La Jolla, CA). Fiji is Just ImageJ (FIJI) was downloaded from https://fiji.sc/. All other materials were purchased from Thermo-Fisher (Toronto, Canada). The CD93 constructs were described previously (28).

### CHO Cell Culture and Transfection

CHO cells were cultured in F-12 media + 10% FBS in a 37 °C/5% CO_2_ incubator. Upon reaching >80% confluency, CHO cells were split 1:10 by washing with PBS, incubating for 2 min at 37°C with 0.25% Trypsin-EDTA, and then suspension in a 4× volume of media + 10% FBS. For microscopy, ~2.5 × 10^5^ cells were plated onto sterile 18 mm round coverslips placed into the wells of a 12-well tissue culture plate. 18 to 24 hrs later the cells were transfected by adding drop-wise to the cells a mixture comprised of 1 µg/well of either human CD93-GFP or human HA-CD93 and 2.25 µl of GenJet reagent suspended in 76μl of serum-free DMEM. Cells were imaged between 18 and 32 hours after transfection.

### 12CA5 Hybridoma Culture, Fab Generation and Fluorescent Labeling

12CA5 hybridoma cells were maintained at 37°C/5% CO_2_ in RPMI + 10% FBS and split 1:5 once maximum cell density was reached. For antibody collection, a 60 ml culture at maximum density was pelleted by centrifugation (300 g, 5 min), resuspended in 60 mL of serum-free hybridoma media, and cultured for 5 days. Cells and media were transferred to centrifuge tubes and centrifuged at 3,000 g, 20 min, 4°C to remove cells and particulates. Antibodies were then concentrated and the medium replaced with 10 mM Tris + 150 mM NaCl (pH 7.4) using a 60 kDa cutoff centrifuge concentrator. Concentrated antibody was diluted to 5 mg/ml total protein and Fab fragments generated by adding 10 mM EDTA and 20 mM Cysteine-HCl to 1 ml of antibody solution, followed by addition of 500 μl of immobilized papain and incubation at 37°C for 18 hours. Papain was then removed by a 1,000 g, 15 min centrifugation. Fab fragments were then purified by separating the protein mixture using FPLC equipped with a Sephacryl S100 column. Aliquots corresponding to the first (Fab fragment) peak were pooled and stored for later use at 4°C. Fab fragments were directly conjugated to Alexa 488 or Cy3 using protein conjugation kits.

### Single-Particle Tracking

SPT was performed using a Leica DMI6000B widefield fluorescent microscope equipped with a 100×/1.40NA objective, supplemental 1.6× in-line magnification, a Photometrics Evolve-512 delta EM-CCD camera mounted using a 0.7× C-mount, a Chroma Sedat Quad filter set and a heated/CO_2_ perfused live-cell piezoelectric stage as described previously (28). Briefly, coverslips were mounted in Leiden chambers and the chambers filled with 1 ml of 37°C imaging buffer (150 mM NaCl, 5 mM KCl, 1 mM MgCl_2_, 100 µM EGTA, 2 mM CaCl_2_, 25 mM HEPES and 1500 g/l NaHCO_3_). Samples were maintained at 37°C/5% CO_2_ on a heated/CO_2_ perfused microscope stage for the duration of the experiment. 30 s duration videos of the basolateral side of the cells were recorded at a frame rate of 10 frames/second, with 5 to 15 cells recorded per coverslip. The resulting videos were cropped to the area containing the cell and exported as TIFF-formatted image stacks. Initial analysis was performed using the Matlab scripts of Jaqaman *et al*. (36) on a dual-Xeon E5-2630 chip workstation running OpenSUSI Linux and Matlab 2014b. Custom-written Matlab scripts were then used to remove any CD93 trajectories with a positional certainty worse than ±25 nm (SNR ⪆ 2.2), and remaining datasets containing > 1,000 trajectories retained for further analysis, leading to rejection of ~40% of trajectories. Our previous studies of CD93 diffusion identified no differences in the diffusion coefficient of CD93 on the basolateral or apical cell surface, a minimal detection of vesicular CD93, using this method (28).

### Immunolabeling of CD93

CD93 was immunolabled such that cross-linking was avoided, at a labeling density which allowed resolution of individual point-spread functions, as per our previous studies (28, 37, 38). Briefly, CD93-GFP transfected CHO cells were cooled to 10°C and blocked for 20 min in F12 media + 4% human serum, then incubated for 20 min with blocking solution containing 1:5,000 anti-human CD93. Samples were washed 3 × 5 min in PBS and blocking buffer containing 1:10,000 diluted Cy3-conjugated Fab secondary antibodies added for an additional 20 min. The samples were washed an additional 3 × 5 min in F12 media + 10% FBS and kept at 10°C until imaged.

### Ensemble-Averaged and Time-Averaged MSDs

The CD93 trajectories in each independent experiment were first classified as either freely diffusing or corralled using TAMSD and MSS analysis, utilizing the software of Jaqaman *et al*. for these calculations (36). Data ensembles were then created by binning the CD93 tracks from all experiments based on the size of the confinement regions determined from this initial classification. Tracks from CD93 in confinement zones with diameters of 100 ± 50, 200 ± 50, 300 ± 50, 400 ± 50, and 500 ± 50 nm, and from freely diffusing CD93 were placed into separate ensembles. The EA-MSD,

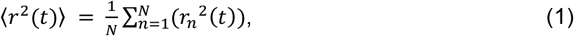

was calculated for each ensemble using a custom-written Matlab script. Here the ensemble average is denoted by the overbar. 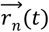 is the position of the *n*^th^ CD93 molecule at experimental time *t*, measured relative to its position at *t* = 0, and *r_n_* is its magnitude. For each particle trajectory, the first frame in which the particle was detected (t’) was defined as t = 0.

A set of trajectories with a given total measurement time *T* was generated by selecting all trajectories within an ensemble having length *T* or greater, then truncating them at the chosen *T* (29). The squared displacement for a given lag time *τ* was calculated for each trajectory, then averaged over all starting times *t’* to give the TA-MSD:

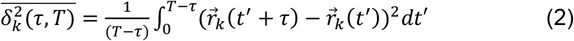

for *τ* ≤ (*T*/3). The subscript *k* labels the trajectory and the angle brackets denote the time average. This quantity was then averaged over all trajectories in the ensemble, yielding the ensemble-averaged, time-averaged MSD, EA-TAMSD 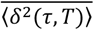:

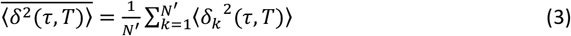

which is a function of both *T* and *τ*. Here the sum is over the *N*‘ truncated trajectories in the ensemble having length *T*.

The time-averaged MSD can be reasonably well described by a power law dependence on the lag time *τ*,

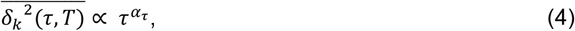

over a range of lag times. As described below, the distribution of the resulting power law exponents α_*τ*_ at a given value of *T* was fitted to a sum of two Gaussians using a maximum likelihood estimation method.

### Mean Maximal Excursion

The above statistics are based on the full distribution of particle displacements. The moments of the distribution of maximal displacements of the trajectories from their starting points can also be used to distinguish different types of anomalous diffusion (39). We first determined *r_M_(t)*, the maximum displacement of each tracked CD93 molecule from its initial position at *t* = 0 over a measurement time *t*. The MME is the average of *r_M_* over an ensemble of particles. The second and fourth moments of the distribution of maximal displacements, referred to as MME moments, are given by

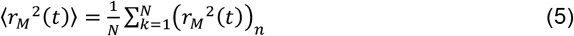

and

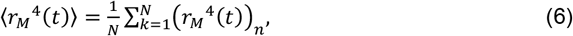

where the overbar indicates an average over the ensemble of *N* trajectories. The ratio

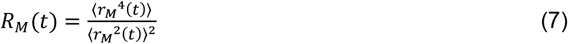

is referred to as the MME ratio. The analogous ratio for the full distribution of displacements,

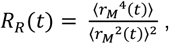

where 〈*r_M_*^2^ (*t*)〉 is the EA-MSD introduced above, and

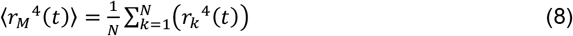

is the fourth moment of the distribution of *r*, is referred to as the regular ratio.

The values of these ratios are different for different models of diffusion. For Brownian motion, *R_R_* = 2 and *R_M_* = 1.49. Diffusion on a fractal gives *R_R_* < 2 and *R_M_* < 1.49 while fractional Brownian motion gives *R_R_* = 2 and *R_M_* < 1.49. Values of these ratios higher than those for normal Brownian motion indicate diffusion by a CTRW (39).

### Windowed Diffusion Analysis

To assess the temporal evolution of diffusion in individual CD93 trajectories over the course of the experiment, in real-time, the ensemble of trajectories for all corralled CD93 molecules was filtered to remove any trajectories less than 5 s in duration. A window encompassing the current time point ± 5 frames was applied to each point of each track, the average particle position calculated for each window, and MSS analysis (36, 40) applied to the segment of the trajectory within the window. Based on the results of this analysis as a function of time over the length of the trajectory, the corralled tracks could be classified as stably corralled (for which all windows were classified as corralled, with no significant change in the average particle position); undergoing hop diffusion (all segments classified as corralled, but with more than one distinct particle position with no overlap of the corral area); undergoing transient escape (initially corralled tracks containing periods of free diffusion separated by periods of corralling, or tracks which escaped corralling within 2 s of the end of the track), or escaped (free diffusion for ≥2 s without re-confinement for the remainder of the track).

### Surface Confinement and Endocytosis Analysis

Surface CD93 was labeled at saturation by incubating with a 1:100 dilution of a 1:10,000 mixture of unlabeled:Cy3-labeled 12CA5 Fab for 10 min at 10°C. Cells were then washed 3× with PBS and stored at 10°C until imaged, at which point they were mounted in Leiden chambers filled with 37°C imaging buffer containing 1:1,000 dilution of Alexa-488 conjugated 12CA5 Fab fragments. For diffusion analysis, five random fields were selected for imaging and short (50 frame) 488 and Cy3 channel SPT videos were collected in each field every 5 min for 30 min. Cy3-labeled and 488-labeled CD93 were individually tracked and classified as free or corralled as described above. As controls, cells fixed for 20 min with 4% PFA prior to labeling, or live cells pre-treated for 20 minutes with 1mM N-ethylmaleimide or 5µg/ml brefledin A, were labeled as above and the number of 488-labeled CD93 present on each cell quantified from images taken every 5 minutes for 30 minutes. To assess colocalization with Arf6 and Rab11, cells ectopically expressing HA-CD93 plus either Arf6-GFP or Rab11-GFP were labeled with saturating concentrations (1:100 dilution) of Cy-3 labeled 12CA5 Fab fragments at 10°C. The cells were then washed 3× with PBS and a 1:1,000 dilution of 647-labeled 12CA5 Fab fragments added to label any CD93 exocytosed during the experiment, and the cells warmed to 37°C for 1 hr. The cells where then fixed with PFA and imaged as described above. The fraction of intracellular CD93 puncta co-localizing with wither Arf6 or Rab11 was then determined using FIJI.

### Ground-State Depletion Microscopy

CD93-expressing CHO cells were treated with Brefeldin A or vehicle for 60 min and fixed with 4% PFA for 20 min at 37°C; fixation was conducted at physiological temperatures to prevent changes in CD93 distribution driven by membrane phase changes. CD93 was then immunolabeled as described above and a second fixation performed with 2% PFA for 10 min at room temperature. The samples were then mounted on a depression slide filled with PBS + 100 mM cysteamine and the coverslip sealed to the slide with VALAP. The sample was immediately transferred to a Leica SR Ground State Depletion microscope and the basolateral membrane imaged in TIRF mode. Using our MIiSR software (38) the resulting images were filtered to remove any CD93 molecules detected with > 20 nm precision, and the filtered images segmented using the OPTICS algorithm. The roundness (41) of each segmented CD93 cluster was calculated as:

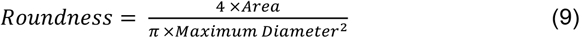

Puncta with a Roundness > 0.5 (for an ellipse, equivalent to a length:width ratio < 2:1) were defined as “Point-Like”, with any cluster not meeting this criteria classified as “Web-Like”

### Statistical Analyses

Unless otherwise noted, data are presented as mean ± SEM and analyzed using ANOVA with Tukey correction in Graphpad Prism. Statistical significance was set at p < 0.05.

## Results

### CD93 Undergoes Anomalous Diffusion

The diffusion of CD93 was recorded using single particle tracking microscopy (SPT), CD93 trajectories reconstructed using a robust tracking algorithm which utilizes mixture model fitting to detect single fluorophores in dense fluorophore fields and a global combinatorial optimization approach to link particles from successive frames into tracks (36), followed by MSS analysis applied to TAMSDs calculated for individual tracks (40) to identify freely diffusing and corralled populations of CD93. As is typical of transmembrane proteins, a large fraction of CD93 underwent corralled diffusion (Fig. 1A), indicative of trapping in membrane corrals (37, 42–45). These corrals ranged in size, averaging 192 nm in diameter (Fig. 1B). Using the conventional method of quantifying confined diffusion coefficients by calculating a TAMSD, at short lag times (τ < 0.5 s), we determined that diffusion occurs more slowly in corrals (Fig. 1C). However, this analysis must be treated with caution as it assumes a linear dependence of TAMSD on lag time, while MSS analysis identifies corralled populations based on a non-linear dependence (40). Moreover, this analysis uses only lag-times, potentially obscuring more subtle anomalous diffusion behavior (29).

**Figure 1:**
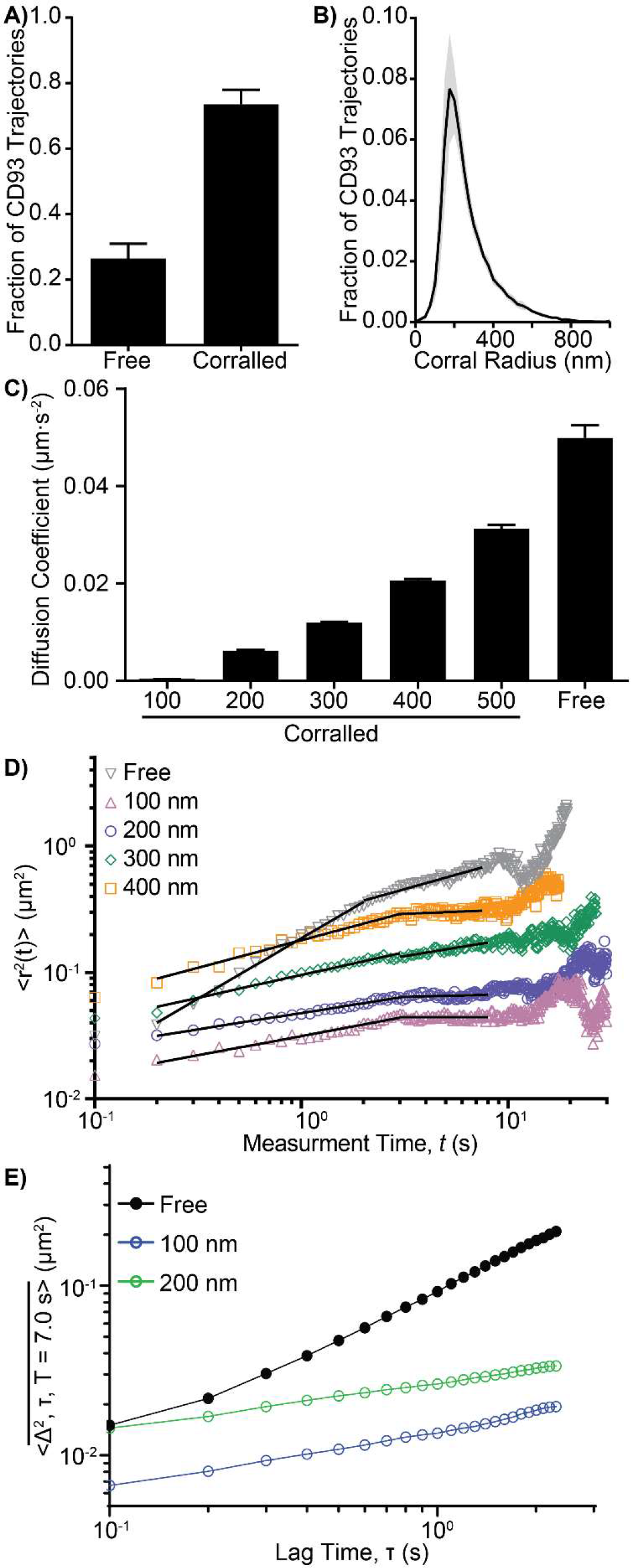
CD93 Diffusion is Non-Brownian. The diffusion of CD93-GFP ectopically expressed on CHO cells was studied by single particle tracking. A) Portion of CD93 undergoing corralled versus free diffusion. B) Distribution of CD93 confinement zone sizes. C) Diffusion coefficients, determined using lag times of less than 100 ms, of freely diffusing CD93 and CD93 corralled in corrals of 100 nm to 500 nm diameter. D) Effect of time on the ensemble-averaged MSD of CD93-GFP. E) The effect of lag time on the EA-TAMSD of freely diffusing CD93 and CD93 in 100 nm and 200 nm corrals. n = mean ± SEM of a minimum of 3 independent experiments (A-C) or the mean of a track ensemble containing 31,633 trajectories collected over 3 experiments. * = p < 0.05, Paired t-test (A) or ANOVA with Tukey correction (B-C), compared to freely diffusing CD93

We reanalyzed CD93 diffusion using an ensemble-averaged MSD (EA-MSD, 〈*r_M_*^2^(*t*)〉), as the behavior of an EA-MSD model can help identify the correct diffusion model. Unlike TAMSD analyses, EA-MSD analysis calculates diffusion coefficients from particle displacements measured relative to the initial particle position, using measurement time *t* rather than lag time τ, therefore allowing for the identification of processes which evolve over the course of the experiment. Unexpectedly, the EAMSDs for both free and corralled CD93 displayed a biphasic dependence on *t*, with 〈*r_M_*^2^(*t*)〉 initially increasing, followed by a plateau after ~2 s of experimental time (Fig. 1D). For corralled populations this plateau is consistent with confinement in both Brownian and non-Brownian models of diffusion, but it is unexpected for a freely diffusing population (6, 30, 32). The power-law indices for the rising (α_1_) and plateau (α_2_) segments of each EA-MSD plot were calculated (Table 1). As expected, both α_1_ and α_2_ were less than 1 for corralled CD93. For CD93 initially classified as freely diffusing, the EA-MSD was linear at low values of t (α_1_ ≈ 1.0), consistent with Brownian motion. However, for *t* > 2.0 s, the motion of these molecules became subdiffusive, with α_2_ ≈ 0.46. This behavior is inconsistent with Brownian diffusion and indicates that even those molecules initially classified as freely diffusing are in fact undergoing some form of anomalous diffusion. Next, the ensemble-averaged time-average MSDs (EA-TAMSD, 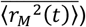 were calculated for freely diffusing CD93 and CD93 in 100 nm and 200 nm corrals (Fig. 1E). The nearly linear relationship between EA-TAMSD and *τ* observed for the free CD93 ensemble is inconsistent with fractional Brownian motion or motion in a fractal environment, while the sub-linear dependence of the EA-TAMSD on *τ* observed for confined CD93 is consistent with several models of anomalous diffusion.

**Table 1:** Power-Law Indices of EA-MSD Plots at Early (0.1–2 s, α_1_) and Late (2 – 12 s, α_2_) Periods of Experimental Time.

| Ensemble | $\alpha_1$ | $\alpha_2$ |
| --- | --- | --- |
| 100 nm | 0.31 | 0.00 |
| 200 nm | 0.25 | 0.03 |
| 300 nm | 0.36 | 0.28 |
| 400 nm | 0.44 | 0.07 |
| Free | 0.96 | 0.46 |

SPT/MSS analysis (40) was used to generate CD93-GFP ensembles for freely diffusing molecules and molecules in confinement zones ranging from 100 nm to 400 nm in diameter, and the EA-MSD plots for CD93 in all ensembles determined. The power-law indices for the resulting EA-MSD plots was then determined at early (0.1 – 2.0 sec) and late (2.0 – 12.0 sec) periods of experimental time.

To investigate this non-Brownian behavior further we determined the power-law index *α_τ_* characterizing the time-averaged MSD (TAMSD,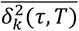) as a function of lag time τ for trajectories of a given length *T*, as in Eq. (4) in the methods. For a population undergoing Brownian diffusion, the statistical distribution of *α_τ_* is expected to be narrow (32). Fig. 2A illustrates the TAMSD for 40 randomly selected CD93 molecules classified as freely diffusing by MSS analysis, for trajectories of length *T* = 2.0 s. In contrast for what is expected for a Brownian population, these molecules showed a high variation in their effective diffusion coefficient. Similar results were obtained for corralled molecules for a range of track lengths and corral sizes, as illustrated in Fig. 2B for 200 nm corrals and *T* = 5.0 s. The distribution of *α_τ_* for the freely diffusing population plotted in Fig. 2A was approximately Gaussian, whereas the distribution for the 200 nm-corralled CD93 plotted in Fig. 2B was distinctly non-Gaussian (Fig 2C). Both the freely diffusing and corralled CD93 had mean power-law indices < 1.0, indicating that both populations were moving subdiffusively.

**Figure 2:**
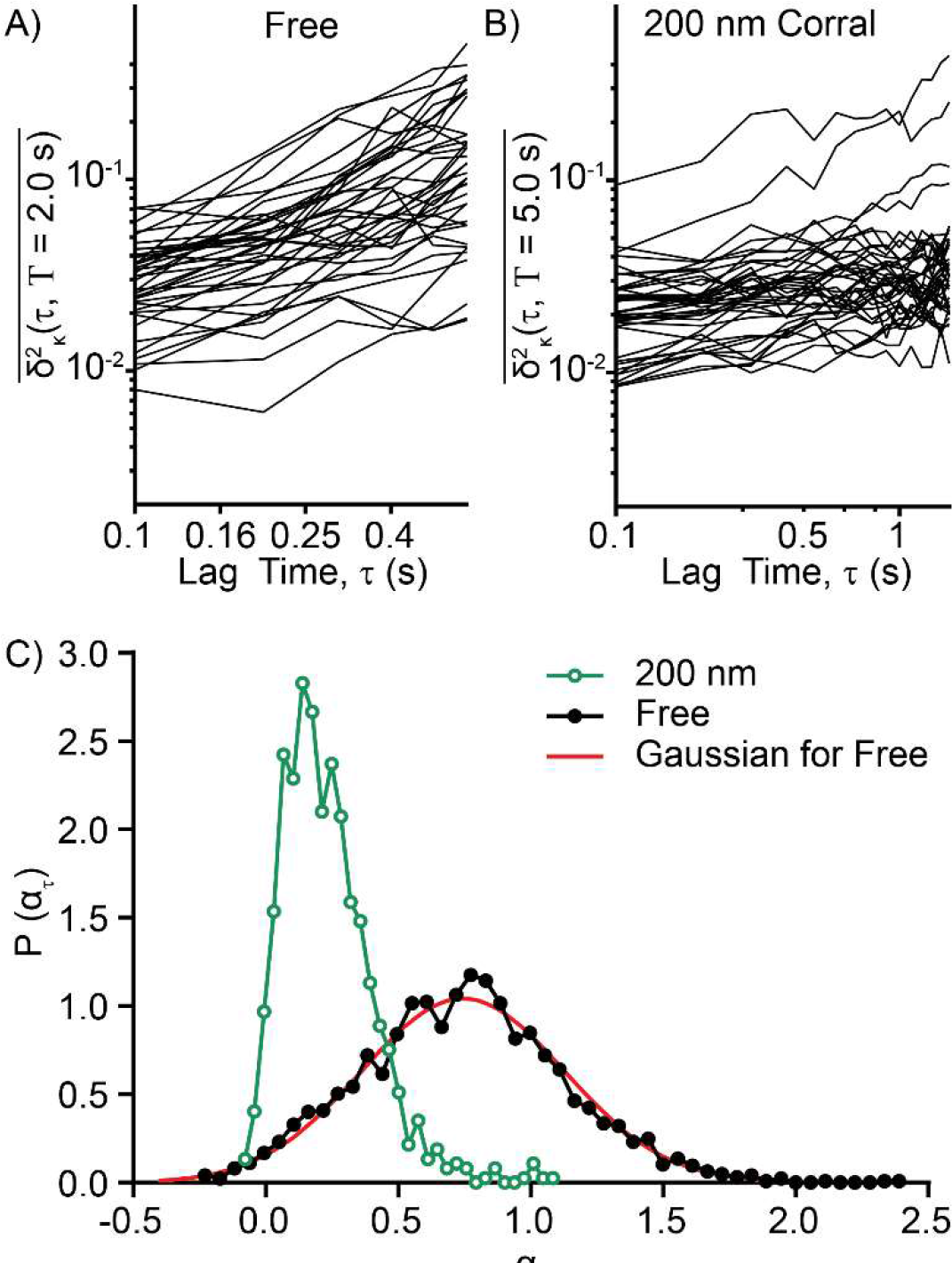
CD93 Diffusional Anomalies Revealed by Time-Averaged MSD. Time averaged MSD curves were calculated for individual CD93-GFP, for trajectory lengths Τ = 2.0 s (free CD93) and 5.0 s (CD93 in 200 nm corrals). A-B) Log-log plots of time-averaged MSD curves for 40 randomly-selected CD93-GFP molecules which were freely diffusing (A) or corralled in 200 nm corrals (B). C) Distribution of the power-law index αT of time-averaged MSD plots for freely diffusing and corralled CD93. The power-law indices of free CD93 were normally distributed with a mean slightly less than the value of 1 expected for Brownian diffusion; αT for CD93 in 200 nm corrals was not normally distributed and had a much smaller mean value. Data are representative of (A-B) or are an ensemble obtained from 3 independent experiments (C).

### CD93 Diffuses by a Continuous Time Random Walk Mechanism

To distinguish between several potential models of anomalous diffusion, we evaluated the dependence of EA-TAMSD on *T* for a lag time of τ = 0.1 s. The lowest available value of *τ* was used to maximize the number of available data points. EA-TAMSD decreases with increasing *T* as a power law with a negative power-law index α_t_ (Fig. 3A). This behavior is consistent with both CTRWs and confined CTRWs, but not with ergodic models of anomalous diffusion such as fractional Brownian motion (29, 32, 46). Table 2 summarizes the power law indices α, α_τ_, and α_t_ that characterize the dependence of EA-MSD on *t*, EA-TAMSD on τ and EA-TAMSD on *T*, respectively. For uncorralled CD93, these indices are in good agreement with the predictions for a subdiffusive continuous time random walk, for which 〈*r*^2^〉 ∝ *t^α^* and 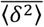 ∝ *τ/T*^1-*α*^, with *α* ≈ 0.5. The indices found for 100 nm and 200 nm corralled CD93 agree well with those expected for particles engaged in a confined, subdiffusive CTRW at long times (*T*≫t, EA-TAMSD ≈ (*τ/T*)^1-*α*^), for which 〈*r*^2^〉 ∝ *t*^0^ and 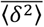 ∝ *τ*⁄/*T*^α^, with *α* ≈ 0.3 (6, 32, 46). To further confirm that CD93 was diffusing via a CTRW, mean-maximal excursion (MME) analysis was applied to the freely diffusing CD93 ensemble and found to be consistent with diffusion via a CTRW for all *t* > 0.1 s (Fig. 3B). Consistent with a CTRW, the regular moment ratio *R_R_* was consistently greater than 2.0, and the MME moment ratio *R_M_* above 1.49, for all evaluated measurement times (Fig. 3C). In contrast, these results are not consistent with FBM or diffusion on a fractal, for which *R_M_* < 1.49, *R_R_* = 2 (FBM) and *R_M_* < 1.49, *R_R_* < 2 (fractal) respectively. Our data therefore indicate that freely diffusing CD93 in fact moves via a subdiffusive CTRW.

**Figure 3:**
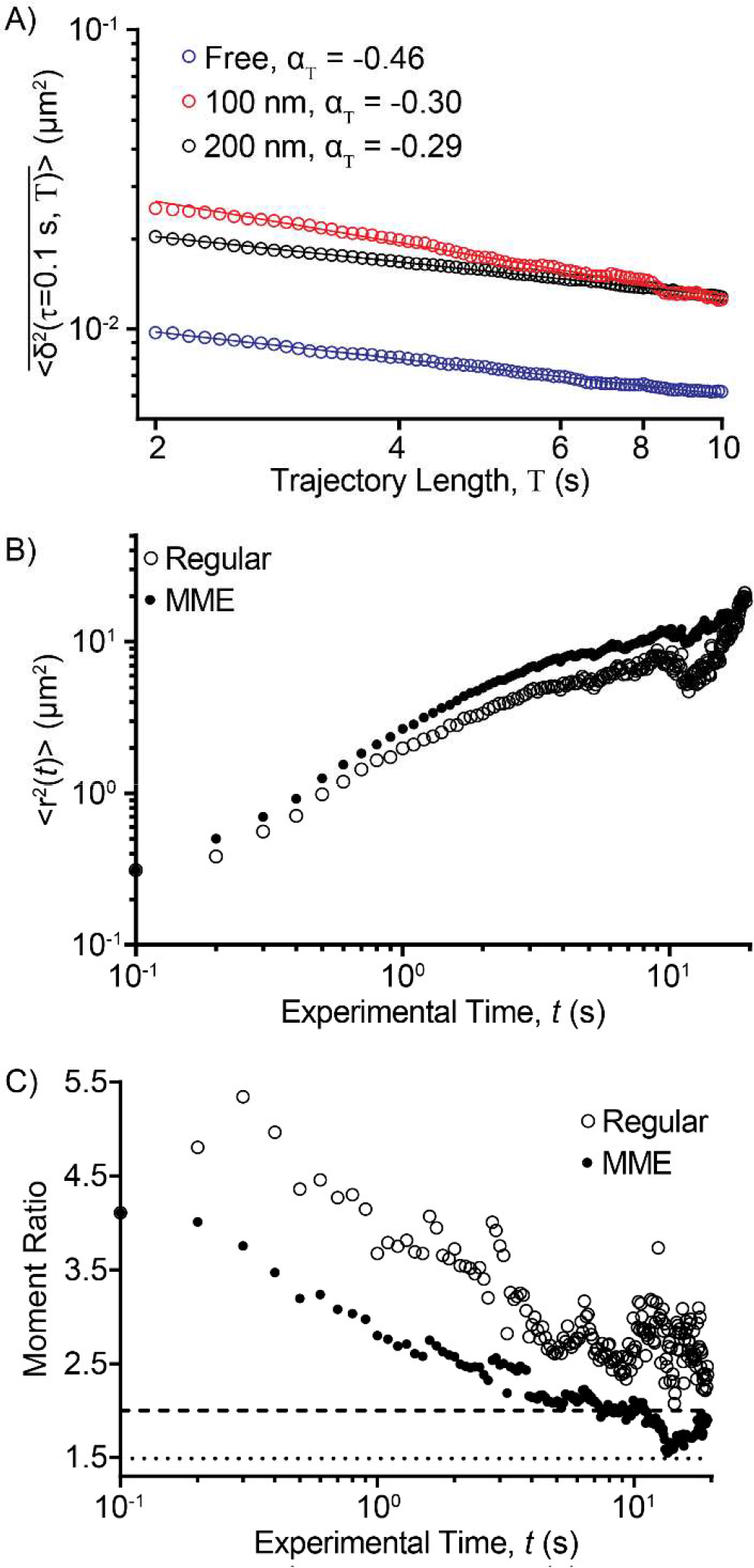
CD93 Diffusion is Consistent with a CTRW. Mean-maximal excursion analysis of freely diffusing CD93 was performed to identify the model that best describes the diffusion of CD93. **A**) Effect of trajectory length on the EA-TAMSD. **B**) The effect of experimental time on the EAMSD (Regular) and second MME (MME) moments. **C**) The effect of experimental time on the ratios between the second and fourth EA-MSD moments (Regular) and the second and fourth MME moments (MME). Horizontal lines indicate the regular (dotted line) and MME (dashed line) moment ratios predicted for Brownian motion; ratios above these values are expected for a CTRW. Data are the ensemble of 3 independent experiments.

**Table 2:**
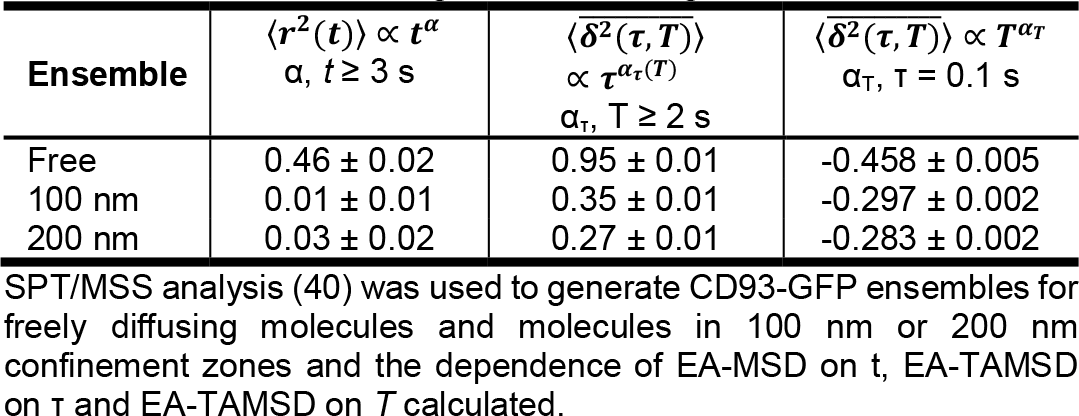
Power-Law Indices of CD93 MSD’s, Ensemble-Averaged MSD’s and Ensemble-Averaged, Time-Averaged MSD’s.

### Non-symmetric Evolution of CD93 Diffusion

To investigate the temporal evolution of CD93 diffusion, we investigate the dependence of *α_τ_* on the trajectory length T for both free and corralled CD93. For corralled CD93, the probability distribution P of *α_τ_* was roughly Gaussian with a mean that did not vary significantly with *T*, indicative of a single population whose diffusional properties do not change over time (Fig. 4A). In marked contrast, P(*α_τ_*) for freely diffusing CD93 is Gaussian with a mean of µ_1_ = 0.756 for short trajectories, while for longer tracks, a second peak centered at μ_2_ = 0.197 emerges (Fig. 4B). This indicates the appearance over time of a second, distinct population of molecules with different diffusion behavior, suggesting that some of the CD93 which was initially freely diffusing becomes trapped in corrals as the experiment progresses (Fig. 4B). To investigate this further, we fitted the P(*α_τ_*) data for each trajectory length to a sum of two Gaussians and extracted the mean value of *α_τ_* for the high-mobility (µ_1_) and low-mobility (µ_2_) populations. Both µ_1_ and µ_2_ were independent of *T* (Fig. 4C). This indicates that the diffusional environment of the two populations did not change significantly over the duration of the experiment. The µ_2_ (corralled) peak was first detectable at *T* ≈ 2.0 s, and the fraction of the detected CD93 contributing to this peak increased approximately linearly with *T*, with 19 ± 6% of the CD93 that was originally freely-diffusing becoming corralled after 8 s of observation (Fig. 4D). This suggests that the diffusion of CD93 evolves in a non-symmetric manner over real-time: freely diffusing CD93 becomes increasingly corralled, without being replenished by CD93 escaping from corrals. In the absence of other replenishment processes, this would inevitably lead to the confinement of all CD93 in corrals. Since this contradicts what is observed, replenishment of free CD93 must occur by another cellular process.

**Figure 4:**
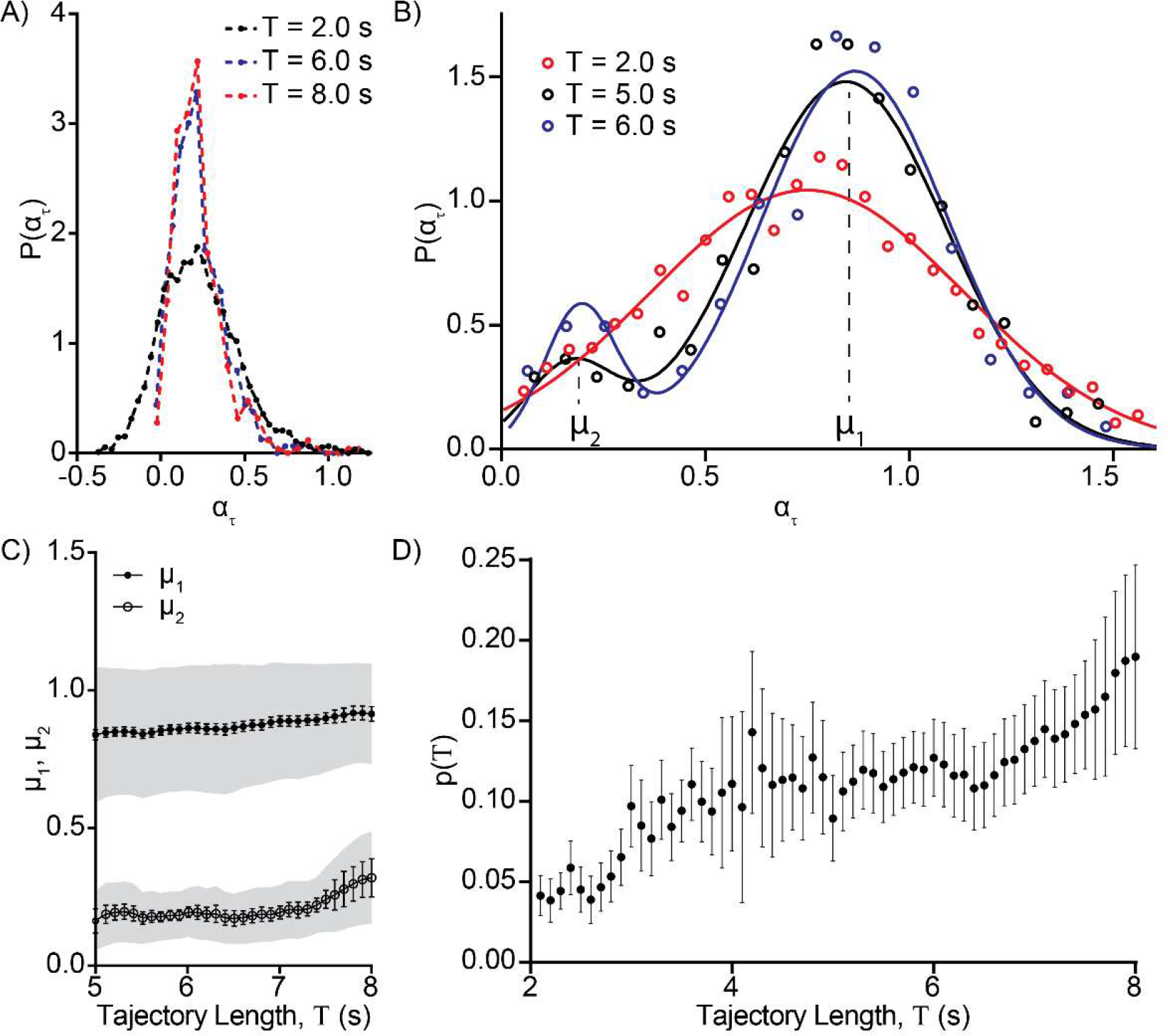
Time-averaged MSDs Evolve Over Experimental Time. Power-law indices were determined from the TA-MSDs of freely diffusing and corralled CD93 for trajectory lengths from 2 to 8 seconds. A) Distribution of power-law indices αT for corralled CD93 at T = 2.0, 6.0 and 8.0 s. B) Distribution of αt for freely diffusing CD93 at δ = 2.0, 5.0 and 6.0 s. The two peaks centered at exponent values μ1 and μ2, indicate the presence of two distinct populations which diffuse at different rates. C) The mean and standard error of αT values for the more diffusive (μ1) and less diffusive (μ2) populations of CD93 originally identified as uncorraled, calculated by fitting the distribution of power-law indices to the sum of two Gaussians for total measurement times from 5 to 8 s. D) The fraction of CD93 initially classified as freely diffusing that appears in the less-diffusive population as a function of T. n = 3, Data are presented as mean ± SEM (error bars) or mean ± SD (shaded areas).

### Endocytosis and Exocytosis Maintain Freely Diffusing CD93 Populations

We confirmed that the observed change in diffusion behavior with track length was not an artefact of our analysis methods by breaking the trajectories of CD93 initially classified as corralled into windows 1 s in duration, and investigating how the diffusional behavior and position of each trajectory changed as the window was progressively moved through the time-course. The vast majority of CD93 was found to be either stably corralled or undergoing hop diffusion, with only 272 of the 311,347 analyzed CD93 trajectories undergoing transient or total escape from corrals (Fig 5A). This confirms that once CD93 is corralled, it remains corralled. The freely diffusing population of CD93 must therefore be replenished by a different process. The endocytic uptake of receptors from the plasma membrane, followed by their replenishment through a combination of exocytosis from recycling compartments and secretion of *de novo* synthesized CD93 from the ER/Golgi, are possible mechanisms by which the freely diffusing population of CD93 could be maintained. To test this, Chinese hamster ovary (CHO) cells were transfected with CD93 bearing a single extracellular HA tag and cell-surface CD93 labeled at saturating levels with a 10,000:1 ratio of unlabeled:Cy3-labeled monoclonal anti-HA Fab fragments. Single particle tracking was then performed in a solution containing Alexa488-labeled anti-HA Fab fragments, to label any CD93 exocytosed over the course of the experiment. We then recorded SPT videos of 488- and Cy3-labeled CD93, imaging the same cells every 5 min over a 60-minute period. The number of Alexa-488 labeled CD93 molecules increased linearly with time. Similar increases were not observed in cells fixed prior to labeling or in cells in which exocytosis was blocked with 1mM N-ethylmaleimide, while blocking Golgi export with brefeldin A had a partial inhibitory effect that strengthened over time (Fig 5B). A much larger portion of recently exocytosed (488-labeled) CD93 underwent free diffusion than did the preexisting surface (Cy3-labeled) CD93, and, in accordance with the above results, the corralling of the recently released CD93 increased over time (Figs. 5C). Interestingly, the portion of preexisting surface (Cy3-labeled) CD93 undergoing free diffusion did not change with time, suggesting that recently synthesized CD93 is not the only source of labeled free CD93 (Fig. 5D). Consistent with this, preexisting surface (Cy3-labeled) CD93 was found to be blebbing, precluding assessment of CD93 distribution in the absence of both endocytosis and exocytosis. However, Brefledin A inhibition of exocytosis from the ER/Golgi depleted approximately 50% of surface CD93, through selective depletion of the punctuate portion of surface CD93, indicating a role for exocytosis in generating the heterogeneous distribution of CD93 in the plasma membrane (Fig 5H-I).

**Figure 5:**
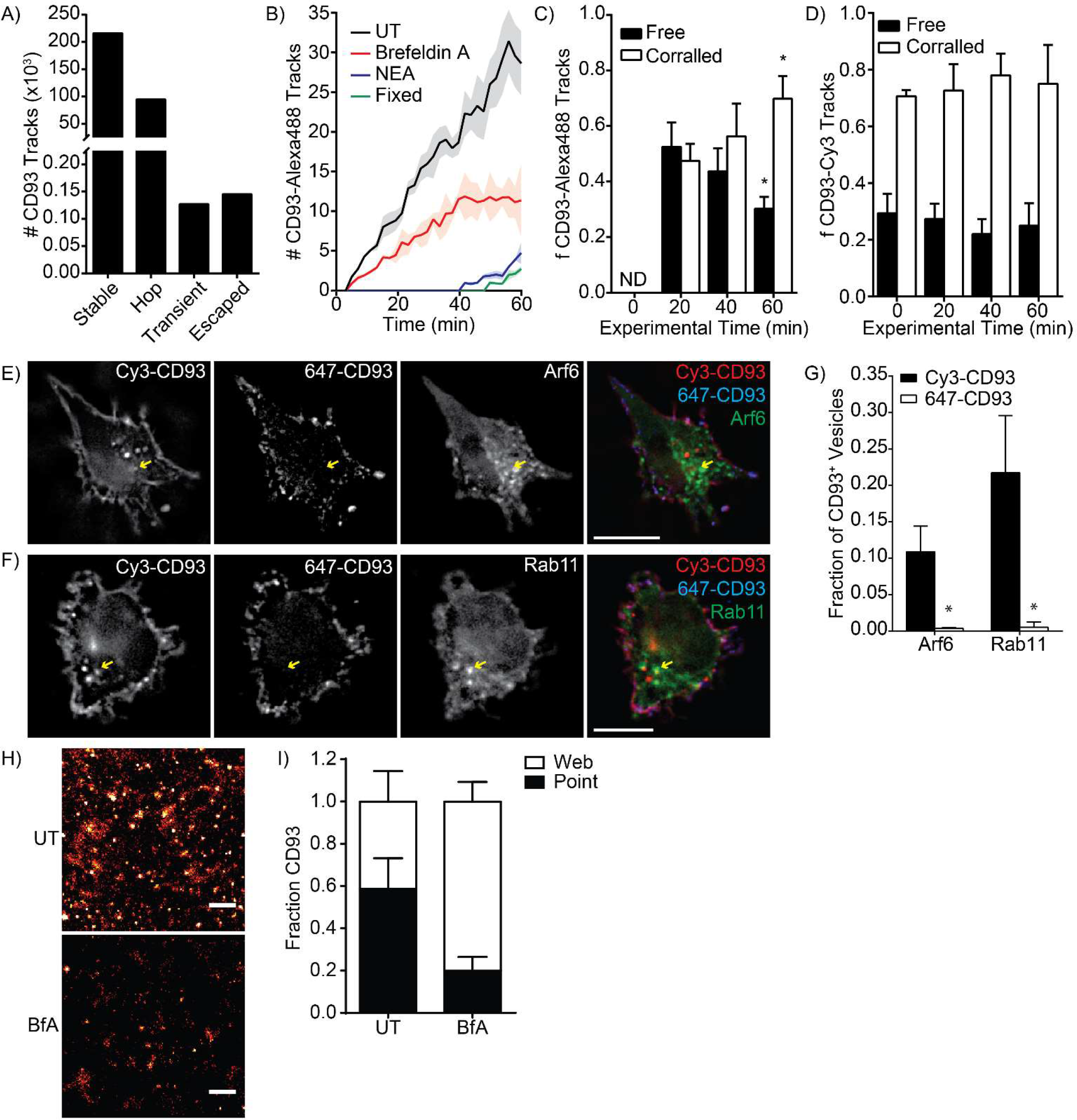
CD93 Diffusion is Sustained by Membrane Recycling. **A)** The number of particle tracks for corralled CD93 classified as stabile corralled (Stable), undergoing hop-diffusion (Hop), transiently escaping confinement (Transient) or totally escaping confinement (Escaped). **B)** Number of newly exocytosed CD93 tracks measured following treatment with Brefeldin A, N-ethylmaleimide (NEA) or fixation of the cell with PFA (Fixed). **C)** Portion of CD93 exocytosed during imaging classified as freely diffusing and corralled, over the course of the experiment. **D)** Portion of CD93 present on the cell surface prior to imaging classified as freely diffusing and corralled, over the course of the experiment **E-G)** Representative images (E-F) and quantification (G) of co-localization of HA-CD93 present on the cell surface at the beginning of the experiment (Cy3-CD93) and recently exocytosed CD93 (647-CD93) with Arf6-GFP or Rab11-GFP after a 1 hr incubation. Arrows indicate vesicles positive for Cy3-CD93 and Arf6 or Rab11. Scale bars are 10 μm. **H)** Super-resolution images of the distribution of CD93 in untreated (UT) versus Brefeldin A (BfA) treated cells. Scale bars are 500 nm. Images are presented as heat-maps. **I)** Effect of Brefeldin A on the fraction of CD93 located in puncta (Point) versus web-like structures (Web). Data are presented as mean (A) or mean ± SEM (B-D, G), n = 5. * = p < 0.05 compared to the same group at 20 min (C-D, ANOVA with Tukey correction) or to Cy3-CD93 in the same cells (G, paired *t*-test). ND = none detected.

## Discussion

Ever since the fluid-mosaic model of the plasma membrane was proposed by Singer and Nicolson (47), efforts have been made to develop a model that accurately describes molecular motion in the plasma membrane. But while early models proposed that membrane diffusion would occur as Brownian diffusion in a 2D planar fluid (48), experiments on intact cells revealed that membrane diffusion is non-Brownian, and moreover, determined that the same proteins and lipids can display different diffusional behavior depending on the temporal resolution and duration over which diffusion is measured (reviewed in 2, 45). This complex diffusional behavior appears to be due to several factors, including molecular crowding within the membrane, the presence of diffusion-limiting cellular structures such as membrane-proximal actin corrals, and heterogeneity in membrane composition (5, 15, 23, 29, 50, 51). More recent work has led to a range of anomalous diffusion models being proposed as possible descriptors of diffusion in cellular membranes, but it remains unclear which, if any, of these models accurately describes plasma membrane diffusion (29, 49). In this study, we have demonstrated that diffusion of the single-pass membrane protein CD93 is well-described by a continuous-time random walk, and specifically eliminated fractional Brownian motion and diffusion on a fractal as possible mechanisms of CD93 diffusion. Moreover, we showed that the trapping and endocytosis of CD93 in membrane corrals, in parallel with exocytosis of CD93 from the ER/Golgi and recycling pathways, are required to maintain this diffusional pattern.

The diffusional behavior of membrane proteins provides information about the underlying molecular and cellular processes producing the behavior, with different molecular and cellular interactions leading to differing relationships between molecular step size and frequency, and to different behavior on different time scales. Brownian motion occurs only when diffusing molecules engage in uncorrelated, elastic collisions in a uniform environment (2). This is not a realistic description of the motion of proteins in biological membranes, given the high molecular packing and the presence of a variety of intermolecular interactions, microdomains, and diffusional barriers that act over a broad range of spatial and temporal scales. FBM, diffusion on a fractal, and CTRWs have all been proposed as descriptors of diffusion in the complex environment of a biological membrane (2, 29, 52). A particle diffusing on a fractal exhibits subdiffusive motion governed by the dimension of the fractal (53). Subdiffusive FBM has been applied to characterize motion in a viscoelastic medium with long time correlation, wherein particles undergoing FBM revisit past locations (32). Temporary immobilization of membrane molecules, conversely, has been described with the aid of continuous time random walks. The CTRW model has previously been applied to characterize the diffusion of Kv2.1 potassium channels in the plasma membrane and the motion of water molecules on the membrane surface (29, 54). Our initial statistical analysis of CD93 motion with respect to the dependence of the EA-MSD on time revealed that CD93 diffusion is anomalous and subdiffusive, with plateaus characteristic of particle confinement for the corralled CD93 populations. The observed subdiffusive behavior is a feature of several diffusion models, including CTRWs, FBM and diffusion on a fractal (32). The high variation in the diffusion coefficients of the TAMSDs of individual CD93 tracks is, however, more consistent with the CTRW model than with FBM (6, 30). Likewise, the nearly-linear relationship of the EA-TAMSD with *τ* observed for free CD93 is consistent with the CTRW model, but not with fractional Brownian motion or motion in a fractal environment (32). Furthermore, the dependence of the EATAMSD on trajectory length observed for all CD93 populations, indicative of system ageing, is in line with the behavior predicted for CTRWs but not for ergodic processes such as FBM or motion in a fractal environment. Overall, the properties and power-law indices of freely diffusing CD93 were found to be consistent with a subdiffusive CTRW, while corralled CD93 was consistent with a confined subdiffusive CTRW. Indeed, CD93 trajectories defined as freely diffusing by the MSS software (36, 40) have EA-MSD and EA-TAMSD characterized by 〈*r*^2^〉 ∝ *t^α^* and 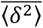∝ (*τ/T*^1-*α*^) respectively, with *α* ≈ 0.5, in good agreement with the predictions for subdiffusive continuous time random walks (32, 46). Mean maximal excursion analysis supported this result (39). CD93 defined as corralled by MSS analysis (MSD power-law index *α* < 1.0) showed different relationships between EA-MSD, EA-TAMSD and t, t and *T*, with 〈*r*^2^〉 ∝ *t*^0^ and 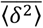 ∝ (*τ/T*)^*α*^, for *α* ≈ 0.3, consistent with diffusion via a confined subdiffusive CTRW occurring over long time scales (6, 30).

Subdiffusive CTRWs arise when particles move by a series of random jumps interrupted by “sticking” or “trapping” events with power-law distributed waiting times. The immobilization events underlying CD93 motion in the model of a CTRW may have several biological origins. In the plasma membrane, temporary molecular immobilization is thought to arise from transient interactions with a macromolecular complex or a cytoskeletal component (2). Most interactions possess a specific dissociation coefficient, which results in an exponentially-distributed waiting-time distribution (29). Power-law distributed waiting times can, however, arise from non-stationary or heterogeneous interactions. For example, if a particle binds to a growing macromolecular complex its dissociation coefficient may change with time. In cases where the probability of its escape decreases as the complex grows, the particle dissociation times have been shown to become power-law distributed, resulting in an evolution towards a CTRW over time (29, 55). Alternatively, a CTRW may arise if a particle interacts with more than one binding partner or macromolecular complex, resulting in a distribution of dissociation coefficients that leads to a heavy-tailed power-law distribution of waiting times. Nothing precludes such interactions from occurring for CD93, either within or outside of confinement zones. Indeed, CD93 has been reported to interact with both moesin and GIPC through its cytosolic tail, as well as with cholesterol-stabilized membrane cages independently of its intracellular domain (56, 57). In principle, these interactions could result in the power-law distributed waiting times characteristic of a CTRW, and are to be investigated further. While these studies and our data are consistent with non-ergodic diffusion occurring via a CTRW driven by interactions with macromolecular or cytoskeletal structures, other studies have ascribed this breaking of ergodicity by membrane proteins to diffusion in a heterogeneous diffusion environment (50, 51). Both mechanisms likely contribute to the observed diffusion patterns, potentially operating at different length and time scales. Indeed, transient structures such as lipid rafts (<20 nm, lifespans of 10-20 ms) would create a heterogeneous diffusive environment not observable at the time- and spatial-scales of our experiments, but would account for the subdiffusive behaviors observed and modeled by others at these spatiotemporal scales (8, 13). Additional sources of nonergodicity may include cellular processes and structures – for example, membrane flow created by exocytosis and endocytosis, and actin barriers which form compartmentalized “corrals” in the plasma membrane (16, 58–61). The system ageing associated with CTRWs implies the eventual caging of all freely diffusing particles, unless the pool of freely diffusing particles is somehow replenished. Indeed, in this study we directly observed a gradual reduction in the diffusion coefficient of free CD93. The short-term reduction in the diffusion coefficient of free CD93 over time was found to be due to their motion by a subdiffusive CTRW. The reduction in the diffusion coefficient of free CD93 at longer measurement times (> 8 s) is, however, due to another biological process – the capture of freely diffusing CD93 into membrane corrals. The capture of freely diffusing molecules into membrane corrals has been previously reported for several proteins, and is believed to be a major component of hop diffusion within cellular membranes (4, 5, 15). Contrary to the hop diffusion model of membrane diffusion, in which trapping in corrals is generally on the scale of milliseconds, we observed this trapping to be stable, with corralled CD93 rarely escaping from corrals over periods of many seconds. Indeed, our data suggest that corralling is a prelude to endocytosis – with the endocytosis of corralled CD93 and subsequent exocytosis of *de novo* synthesized and recycled CD93 maintaining the freely diffusing population of CD93. Consistent with our observations, it has been shown that sites of endocytosis restrict diffusion, and that actin (and therefore actin-dependent corrals) must be cleared from secretory vesicle fusion sites prior to exocytosis (62–64). The colocalization of CD93 with the Rab11 and Arf6 recycling compartments, and the observation that blocking Golgi export only blocked ~50% of CD93 exocytosis, indicates the importance of both recycling and *de novo* CD93 synthesis for maintaining normal CD93 diffusion. This is likely a process ubiquitous for most cell-surface receptors – indeed, several receptors, including phagocytic Fc receptors and the endocytic receptor CD36 are known to require on-going recycling and maintenance of an appropriate diffusional environment for their activity. This is consistent with processes analogous to the caging-endocytosis-exocytosis process we have observed here being required for the maintenance of proteins other than CD93 (26, 42, 65–67). Moreover, our data demonstrate that CD93 is non-uniformly distributed in the plasma membrane, with exocytosis from the ER/Golgi responsible for maintaining highly punctuate CD93 clusters in the plasma membrane. These data are consistent with the model proposed by Gheber *et al*., which predicted that the delivery and removal of membrane proteins by vesicular trafficking would be required to form and maintain stable membrane domains (68), and demonstrated that both vesicular trafficking and diffusion barriers determine the size and lifetime of protein clusters on the cell surface (58–60).

In summary, we have demonstrated that the type 1 membrane protein CD93 diffuses in the plasma membrane via a continuous-time random walk mechanism or by a similar non-ergodic diffusive process, and that the aging and continued function of this CTRW process is dependent on a sequential process comprised of the trapping of freely diffusing CD93 into corrals, endocytosis of corralled CD93, and regeneration of the freely diffusing pool of CD93 by the exocytosis of both *de novo* synthesized and recycled CD93. While our investigation was limited to CD93, the motion of other membrane proteins – including the multi-span Kv2.1 potassium channel – has been ascribed to a CTRW (29), suggesting that these processes regulating CD93 diffusion identified in this study may represent a general model of protein diffusion in the plasma membrane.

## Author Contributions

MG performed all experiments except those in Figure 5. MG and JRdB developed the analysis framework for diffusion analysis. BH performed the experiments in Figure 5 and oversaw the study. All authors contributed to design and analysis of the experiments and to the writing of this manuscript.

## Acknowledgements

The authors would like to thank Jessica Ellins for generating the CD93-GFP and HA-CD93 constructs. This study was funded by National Sciences and Engineering Research Council of Canada Discovery Grants to BH and JRdB and a Canadian Institutes of Health Research Operating Grant (MOP-123419) to BH. MG was funded by a NSERC postgraduate scholarship. The funding agencies had no role in study design, data collection and analysis, decision to publish, or preparation of the manuscript.

## References

1. Kusumi, A., C. Nakada, K. Ritchie, K. Murase, K. Suzuki, H. Murakoshi, R.S. Kasai, J. Kondo, and T. Fujiwara. 2005. Paradigm shift of the plasma membrane concept from the two-dimensional continuum fluid to the partitioned fluid: high-speed single-molecule tracking of membrane molecules. Annu. Rev. Biophys. Biomol. Struct. 34: 351–78.

2. Krapf, D. 2015. Mechanisms underlying anomalous diffusion in the plasma membrane. Curr. Top. Membr. 75: 167–207.

3. Wieser, S., M. Moertelmaier, E. Fuertbauer, H. Stockinger, and G.J. Schütz. 2007. (Un)confined diffusion of CD59 in the plasma membrane determined by high-resolution single molecule microscopy. Biophys. J. 92: 3719–28.

4. Fujiwara, T., K. Ritchie, H. Murakoshi, K. Jacobson, and A. Kusumi. 2002. Phospholipids undergo hop diffusion in compartmentalized cell membrane. J. Cell Biol. 157: 1071–81.

5. Murase, K., T. Fujiwara, Y. Umemura, K. Suzuki, R. Iino, H. Yamashita, M. Saito, H. Murakoshi, K. Ritchie, and A. Kusumi. 2004. Ultrafine membrane compartments for molecular diffusion as revealed by single molecule techniques. Biophys. J. 86: 4075–93.

6. Neusius, T., I.M. Sokolov, and J.C. Smith. 2009. Subdiffusion in time-averaged, confined random walks. Phys. Rev. E. Stat. Nonlin. Soft Matter Phys. 80: 11109.

7. Bel, G., and E. Barkai. 2005. Weak Ergodicity Breaking in the Continuous-Time Random Walk. Phys. Rev. Lett. 94: 240602.

8. Eggeling, C., C. Ringemann, R. Medda, G. Schwarzmann, K. Sandhoff, S. Polyakova, V.N. Belov, B. Hein, C. von Middendorff, A. Schönle, and S.W. Hell. 2009. Direct observation of the nanoscale dynamics of membrane lipids in a living cell. Nature. 457: 1159–1162.

9. Hammond, G.R. V, Y. Sim, L. Lagnado, and R.F. Irvine. 2009. Reversible binding and rapid diffusion of proteins in complex with inositol lipids serves to coordinate free movement with spatial information. J. Cell Biol. 184: 297–308.

10. Munguira, I., I. Casuso, H. Takahashi, F. Rico, A. Miyagi, M. Chami, and S. Scheuring. 2016. Glasslike Membrane Protein Diffusion in a Crowded Membrane. ACS Nano. 10: 2584–90.

11. Sonnleitner, A., G.J. Schütz, and T. Schmidt. 1999. Free Brownian motion of individual lipid molecules in biomembranes. Biophys. J. 77: 2638–42.

12. Vaz, W.L., M. Criado, V.M. Madeira, G. Schoellmann, and T.M. Jovin. 1982. Size dependence of the translational diffusion of large integral membrane proteins in liquid-crystalline phase lipid bilayers. A study using fluorescence recovery after photobleaching. Biochemistry. 21: 5608–12.

13. Jeon, J.-H., M. Javanainen, H. Martinez-Seara, R. Metzler, and I. Vattulainen. 2016. Protein Crowding in Lipid Bilayers Gives Rise to Non-Gaussian Anomalous Lateral Diffusion of Phospholipids and Proteins. Phys. Rev. X. 6: 21006.

14. Kusumi, A., T.K. Fujiwara, R. Chadda, M. Xie, T. a Tsunoyama, Z. Kalay, R.S. Kasai, and K.G.N. Suzuki. 2012. Dynamic organizing principles of the plasma membrane that regulate signal transduction: commemorating the fortieth anniversary of Singer and Nicolson’s fluid-mosaic model. Annu. Rev. Cell Dev. Biol. 28: 215–50.

15. Fujiwara, T.K., K. Iwasawa, Z. Kalay, T.A. Tsunoyama, Y. Watanabe, Y.M. Umemura, H. Murakoshi, K.G.N. Suzuki, Y.L. Nemoto, N. Morone, and A. Kusumi. 2016. Confined diffusion of transmembrane proteins and lipids induced by the same actin meshwork lining the plasma membrane. Mol. Biol. Cell. : 11–81.

16. Sadegh, S., J.L. Higgins, P.C. Mannion, M.M. Tamkun, and D. Krapf. Plasma Membrane is Compartmentalized by a Self-Similar Cortical Actin Meshwork. Phys. Rev. X. 7.

17. Sheetz, M.P., M. Schindler, and D.E. Koppel. 1980. Lateral mobility of integral membrane proteins is increased in spherocytic erythrocytes. Nature. 285: 510–1.

18. Schmidt, K., and B.J. Nichols. 2004. A barrier to lateral diffusion in the cleavage furrow of dividing mammalian cells. Curr. Biol. 14: 1002–1006.

19. Saka, S.K., A. Honigmann, C. Eggeling, S.W. Hell, T. Lang, and S.O. Rizzoli. 2014. Multi-protein assemblies underlie the mesoscale organization of the plasma membrane. Nat. Commun. 5: 4509.

20. Espenel, C., E. Margeat, P. Dosset, C. Arduise, C. Le Grimellec, C. a Royer, C. Boucheix, E. Rubinstein, and P.-E. Milhiet. 2008. Single-molecule analysis of CD9 dynamics and partitioning reveals multiple modes of interaction in the tetraspanin web. J. Cell Biol. 182: 765–76.

21. Möckl, L., A.K. Horst, K. Kolbe, T.K. Lindhorst, and C. Bräuchle. 2015. Microdomain Formation Controls Spatiotemporal Dynamics of Cell-Surface Glycoproteins. Chembiochem. 16: 2023–8.

22. Murakoshi, H., R. Iino, T. Kobayashi, T. Fujiwara, C. Ohshima, A. Yoshimura, and A. Kusumi. 2004. Single-molecule imaging analysis of Ras activation in living cells. Proc. Natl. Acad. Sci. U. S. A. 101: 7317–7322.

23. van Zanten, T.S., J. Gómez, C. Manzo, A. Cambi, J. Buceta, R. Reigada, and M.F. Garcia-Parajo. 2010. Direct mapping of nanoscale compositional connectivity on intact cell membranes. Proc. Natl. Acad. Sci. U. S. A. 107: 15437–15442.

24. Yang, X.H., R. Mirchev, X. Deng, P. Yacono, H.L. Yang, D.E. Golan, and M.E. Hemler. 2012. CD151 restricts the α6 integrin diffusion mode. J. Cell Sci. 125: 1478–87.

25. Watanabe, S., B.R. Rost, M. Camacho-Pérez, M.W. Davis, B. Söhl-Kielczynski, C. Rosenmund, and E.M. Jorgensen. 2013. Ultrafast endocytosis at mouse hippocampal synapses. Nature. 504: 242–7.

26. Cox, D., D.J. Lee, B.M. Dale, J. Calafat, and S. Greenberg. 2000. A Rab11-containing rapidly recycling compartment in macrophages that promotes phagocytosis. Proc. Natl. Acad. Sci. U. S. A. 97: 680–5.

27. Kusumi, A., T.K. Fujiwara, N. Morone, K.J. Yoshida, R. Chadda, M. Xie, R.S. Kasai, and K.G.N. Suzuki. 2012. Membrane mechanisms for signal transduction: the coupling of the meso-scale raft domains to membrane-skeleton-induced compartments and dynamic protein complexes. Semin. Cell Dev. Biol. 23: 126–44.

28. Goiko, M., J.R. de Bruyn, and B. Heit. 2016. Short-Lived Cages Restrict Protein Diffusion in the Plasma Membrane. Sci. Rep. 6: 34987.

29. Weigel, A. V, B. Simon, M.M. Tamkun, and D. Krapf. 2011. Ergodic and nonergodic processes coexist in the plasma membrane as observed by single-molecule tracking. Proc. Natl. Acad. Sci. U. S. A. 108: 6438–6443.

30. Burov, S., J.-H. Jeon, R. Metzler, and E. Barkai. 2011. Single particle tracking in systems showing anomalous diffusion: the role of weak ergodicity breaking. Phys. Chem. Chem. Phys. 13: 1800–12.

31. Otten, M., A. Nandi, D. Arcizet, M. Gorelashvili, B. Lindner, and D. Heinrich. 2012. Local motion analysis reveals impact of the dynamic cytoskeleton on intracellular subdiffusion. Biophys. J. 102: 758–67.

32. Metzler, R., J.-H. Jeon, A.G. Cherstvy, and E. Barkai. 2014. Anomalous diffusion models and their properties: nonstationarity, non-ergodicity, and ageing at the centenary of single particle tracking. Phys. Chem. Chem. Phys. 16: 24128–64.

33. Norsworthy, P.J., L. Fossati-Jimack, J. Cortes-Hernandez, P.R. Taylor, A.E. Bygrave, R.D. Thompson, S. Nourshargh, M.J. Walport, and M. Botto. 2004. Murine CD93 (C1qRp) contributes to the removal of apoptotic cells in vivo but is not required for C1q-mediated enhancement of phagocytosis. J. Immunol. 172: 3406–14.

34. Kao, Y.-C., S.-J. Jiang, W.-A. Pan, K.-C. Wang, P.-K. Chen, H.-J. Wei, W.-S. Chen, B.-I. Chang, G.-Y. Shi, and H.-L. Wu. 2012. The epidermal growth factor-like domain of CD93 is a potent angiogenic factor. PLoS One. 7: e51647.

35. Galvagni, F., F. Nardi, M. Maida, G. Bernardini, S. Vannuccini, F. Petraglia, A. Santucci, and M. Orlandini. 2016. CD93 and dystroglycan cooperation in human endothelial cell adhesion and migration adhesion and migration. Oncotarget. 7.

36. Jaqaman, K., D. Loerke, M. Mettlen, H. Kuwata, S. Grinstein, S.L. Schmid, and G. Danuser. 2008. Robust single-particle tracking in live-cell time-lapse sequences. Nat. Methods. 5: 695–702.

37. Heit, B., H. Kim, G. Cosío, D. Castaño, R. Collins, C.A. Lowell, K.C. Kain, W.S. Trimble, and S. Grinstein. 2013. Multimolecular signaling complexes enable Syk-mediated signaling of CD36 internalization. Dev. Cell. 24: 372–83.

38. Caetano, F.A., B.S. Dirk, J.H.K. Tam, P.C. Cavanagh, M. Goiko, S.S.G. Ferguson, S.H. Pasternak, J.D. Dikeakos, J.R. de Bruyn, and B. Heit. 2015. MIiSR: Molecular Interactions in Super-Resolution Imaging Enables the Analysis of Protein Interactions, Dynamics and Formation of Multi-protein Structures. PLoS Comput. Biol. 11: e1004634.

39. Tejedor, V., O. Bénichou, R. Voituriez, R. Jungmann, F. Simmel, C. Selhuber-Unkel, L.B. Oddershede, and R. Metzler. 2010. Quantitative analysis of single particle trajectories: mean maximal excursion method. Biophys. J. 98: 1364–72.

40. Ferrari, R., A.J. Manfroi, and W.R. Young. 2001. Strongly and weakly self-similar diffusion. Phys. D Nonlinear Phenom. 154: 111–137.

41. Solomon, C., and T. Breckon. 2010. Fundamentals of Digital Image Processing. Chichester, UK: John Wiley & Sons, Ltd.

42. Jaqaman, K., H. Kuwata, N. Touret, R. Collins, W.S. Trimble, G. Danuser, and S. Grinstein. 2011. Cytoskeletal Control of CD36 Diffusion Promotes Its Receptor and Signaling Function. Cell. 146: 593–606.

43. Wong, H.S., V. Jaumouillé, B. Heit, S.A. Doodnauth, S. Patel, Y.-W. Huang, S. Grinstein, and L.A. Robinson. 2014. Cytoskeletal confinement of CX3CL1 limits its susceptibility to proteolytic cleavage by ADAM10. Mol. Biol. Cell. 25: 3884–99.

44. Flannagan, R.S., R.E. Harrison, C.M. Yip, K. Jaqaman, and S. Grinstein. 2010. Dynamic macrophage “probing” is required for the efficient capture of phagocytic targets. J. Cell Biol. 191: 1205–18.

45. Jaumouillé, V., Y. Farkash, K. Jaqaman, R. Das, C.A. Lowell, and S. Grinstein. 2014. Actin cytoskeleton reorganization by Syk regulates Fcγ receptor responsiveness by increasing its lateral mobility and clustering. Dev. Cell. 29: 534–46.

46. He, Y., S. Burov, R. Metzler, and E. Barkai. 2008. Random time-scale invariant diffusion and transport coefficients. Phys. Rev. Lett. 101: 58101.

47. Singer, S.J., and G.L. Nicolson. 1972. The fluid mosaic model of the structure of cell membranes. Science. 175: 720–31.

48. Saffman, P.G., and M. Delbrück. 1975. Brownian motion in biological membranes. Proc. Natl. Acad. Sci. U. S. A. 72: 3111–3.

49. Espinoza, F. a, M.J. Wester, J.M. Oliver, B.S. Wilson, N.L. Andrews, D.S. Lidke, and S.L. Steinberg. 2012. Insights into cell membrane microdomain organization from live cell single particle tracking of the IgE high affinity receptor FcεRI of mast cells. Bull. Math. Biol. 74: 1857–911.

50. Charalambous, C., G. Muñoz-Gil, A. Celi, M.F. Garcia-Parajo, M. Lewenstein, C. Manzo, and M.A. García-March. 2017. Nonergodic subdiffusion from transient interactions with heterogeneous partners. Phys. Rev. E. 95: 32403.

51. Manzo, C., J.A. Torreno-Pina, P. Massignan, G.J. Lapeyre, M. Lewenstein, and M.F. Garcia Parajo. 2015. Weak Ergodicity Breaking of Receptor Motion in Living Cells Stemming from Random Diffusivity. Phys. Rev. X. 5: 11021.

52. Höfling, F., and T. Franosch. 2013. Anomalous transport in the crowded world of biological cells. Rep. Prog. Phys. 76: 46602.

53. Meroz, Y., I.M. Sokolov, and J. Klafter. 2010. Subdiffusion of mixed origins: when ergodicity and nonergodicity coexist. Phys. Rev. E. Stat. Nonlin. Soft Matter Phys. 81: 10101.

54. Yamamoto, E., T. Akimoto, M. Yasui, and K. Yasuoka. 2014. Origin of subdiffusion of water molecules on cell membrane surfaces. Sci. Rep. 4: 4720.

55. Weigel, A. V, M.M. Tamkun, and D. Krapf. 2013. Quantifying the dynamic interactions between a clathrin-coated pit and cargo molecules. Proc. Natl. Acad. Sci. U. S. A. 110: E4591–600.

56. Zhang, M., S.S. Bohlson, M. Dy, and A.J. Tenner. 2005. Modulated interaction of the ERM protein, moesin, with CD93. Immunology. 115: 63–73.

57. Bohlson, S.S., M. Zhang, C.E. Ortiz, and A.J. Tenner. 2005. CD93 interacts with the PDZ domain-containing adaptor protein GIPC: implications in the modulation of phagocytosis. J. Leukoc. Biol. 77: 80–9.

58. Lavi, Y., M.A. Edidin, and L.A. Gheber. 2007. Dynamic Patches of Membrane Proteins. Biophys. J. 93: L35–L37.

59. Lavi, Y., N. Gov, M. Edidin, and L.A. Gheber. 2012. Lifetime of Major Histocompatibility Complex Class-I Membrane Clusters Is Controlled by the Actin Cytoskeleton. Biophys. J. 102: 1543–1550.

60. Blumenthal, D., M. Edidin, and L.A. Gheber. 2016. Trafficking of MHC molecules to the cell surface creates dynamic protein patches. J. Cell Sci. 129: 3342–50.

61. Bae, H.-B., J.-M. Tadie, S. Jiang, D.W. Park, C.P. Bell, L.C. Thompson, C.B. Peterson, V.J. Thannickal, E. Abraham, and J.W. Zmijewski. 2013. Vitronectin inhibits efferocytosis through interactions with apoptotic cells as well as with macrophages. J. Immunol. 190: 2273–81.

62. Mitchell, T., A. Lo, M.R. Logan, P. Lacy, and G. Eitzen. 2008. Primary granule exocytosis in human neutrophils is regulated by Rac-dependent actin remodeling. Am. J. Physiol. Cell Physiol. 295: C1354–65.

63. Vitale, M.L., A.R. Del Castillo, L. Tchakarov, and J.M. Trifaro. 1991. Cortical filamentous actin disassembly and scinderin redistribution during chromaffin cell stimulation precede exocytosis, a phenomenon not exhibited by gelsolin. J. Cell Biol. 113: 1057–1067.

64. Van Der Schaar, H.M., M.J. Rust, Chen, H. Van Der Ende-Metselaar, J. Wilschut, X. Zhuang, and J.M. Smit. 2008. Dissecting the cell entry pathway of dengue virus by single-particle tracking in living cells. PLoS Pathog. 4.

65. Freeman, S.A., J. Goyette, W. Furuya, E.C. Woods, C.R. Bertozzi, W. Bergmeier, B. Hinz, P.A. van der Merwe, R. Das, and S. Grinstein. 2016. Integrins Form an Expanding Diffusional Barrier that Coordinates Phagocytosis. Cell. 164: 128–40.

66. Treanor, B., D. Depoil, A. Gonzalez-Granja, P. Barral, M. Weber, O. Dushek, A. Bruckbauer, and F.D. Batista. 2010. The Membrane Skeleton Controls Diffusion Dynamics and Signaling through the B Cell Receptor. Immunity. 32: 187–199.

67. Takahashi, S., K. Kubo, S. Waguri, a. Yabashi, H.-W. Shin, Y. Katoh, and K. Nakayama. 2012. Rab11 regulates exocytosis of recycling vesicles at the plasma membrane. J. Cell Sci. 125: 4049–4057.

68. Gheber, L.A., and M. Edidin. 1999. A Model for Membrane Patchiness: Lateral Diffusion in the Presence of Barriers and Vesicle Traffic. Biophys. J. 77: 3163–3175.

